# Study on Spatial-temporal Variation of Atmospheric Pollution in China between 2014 and 2019

**DOI:** 10.1101/2019.12.20.884346

**Authors:** Guo Peng, Umarova Aminat Batalbievna, Luan Yunqi

## Abstract

The study in this paper reveals that the atmospheric contaminants in mainland China is of concentricity in spatial distribution, persistence in temporal distribution and correlation between different parameters. This spatial-temporal variation law plays an important role in improving and addressing the problem with atmospheric environment of given cities and regions by employing focused and pointed measures. In this paper, seven kinds of atmospheric pollution parameters including PM2.5, PM10, AQI, CO, NO2, O3 and SO2 in 370 Chinese cities from 2014 to 2019 are analyzed based on their hourly mass concentration. The spatial-temporal variations of each parameter in each separated year are obtained by using interpolation calculation towards the annual atmospheric pollution parameters. The results show that higher mass concentration (including the highest mass concentration) of PM2.5, AQI, PM10, CO, NO2, SO2 mainly concentrated in Beijing-Tianjin-Hebei region in the northeast of mainland China and its neighboring regions and Xinjiang region in the northwest of mainland China. The spatial variation of PM2.5, AQI, PM10, CO, NO2, SO2 experienced similar trend. Higher mass concentration (including the highest mass concentration) of O3 mainly concentrated in Qinghai and Inner Mongolia in the central north of mainland China and Shandong on the right side of Beijing-Tianjin-Hebei region. The spatial variation of O3 experienced different trend from that of other parameters. PM2.5, AQI, PM10 indicated increase and decrease, followed by increase and decrease again with time, which was a S-shaped change. Almost the same temporal variation happened to PM2.5, AQI, PM10 and to CO, NO2, SO2, which was opposite to O3. The analysis from the perspective of the annual highest mass concentration of PM2.5, AQI, PM10, NO2 and O3 indicates the atmospheric environment of mainland China had not been authentically improved by 2019. The analysis from the perspective of the annual highest mass concentration of CO and SO2 indicates the atmospheric environment of mainland China had been authentically improved by 2019. What should we do immediately is to strengthen environmental governance and address the source of contamination in Beijing-Tianjin-Hebei region in the northeast of China, Xinjiang region in the northwest of China, Qinghai and Inner Mongolia regions in the central north of China.

## Introduction

At present, a great number of cities in China confront severe situation associated with air pollution, which has become an environmental and social problem drawing much concerns of the general public ^**[1-2]**^. In general, the relative humidity of fog is higher than 90%, but the haze would come into being with this index less than 80% and atmospheric particulates increasing. In people’s daily life, the concentration of atmospheric particulates could witness an upward trend due to many industrial, agricultural and architectural practices, such as the emission of contaminants from power plants, the combustion of such biomass as straw ^**[19]**^, the development of real state and the construction of various infrastructures. Among these, PM2.5, a chief culprit, is a kind of so fine atmospheric particulate (aerodynamic diameter is no more than 2.5 μm) that it could easily pass through the nasal cavity and respiratory tract and directly enter bronchi and pulmonary alveoli. Also, it is of great difficulty for our bodies to release them by depending on metabolic process ^**[6]**^. Meanwhile, the large superficial area and strong adsorption capacity of PM2.5 make itself become a mighty carrier to diffuse microorganisms like bacteria and virus ^**[19]**^, organic cyanides, dioxins^**[3]**^, heavy metals^**[3,14,17]**^ and other kinds of carcinogenic substances. Long-term exposure to such substances could increase the risk of getting cancer to a large extent^**[4]**^. Also, regional pollution^[**8.9**]^ induced by CO, NO2, SO2, O3 ^**[10.11]**^ is increasingly remarkable. In some pivotal cities located in the Beijing-Tianjin-Hebei region and its surrounding areas, Xinjiang, Qinghai, Inner Mongolia, developed Yangtze River Delta region and Pearl River Delta region, haze pollution is frequent and continuous and leads to visibility reduction in a long period^[**12-13**]^. The air quality of these cities is not only far worse than that of developed countries in Europe and America, but also worse than that of other regions in China.

## 1 Sample collection and analysis approach

According to the data derived from the Ministry of Ecology and Environment of the People’s Republic of China^**[24]**^, Beijing Municipal Environmental Monitoring Center^**[23]**^, Beijing Bureau of Statistics^[21]^, the hourly atmospheric pollution parameters related to PM2.5, PM10, AQI, CO, NO2, O3, SO2 in 370 Chinese cities from the year 2014 to 2019 were collected in this paper. The analysis approach employed here is to implement Radial Basis Functions (RBF)-based interpolation calculation towards the annual atmospheric pollution parameters of each category through Arcgis. As a consequence, atmospheric pollution parameters in mainland China could be shown in the form of spatial-temporal distribution diagrams. By further statistics, the top ten cities with higher mass concentration of each air pollution parameter per year between 2014 and 2019 were selected out of the 370 cities, and the frequency of their occurrences was also evaluated and analyzed.

## 2. Discussion and analysis

### 2.1 The spatial-temporal variation of PM2.5 in mainland China from 2014 to 2019

From Figure 1, it can be seen that the Beijing-Tianjin-Hebei region in the northeast of mainland China and Xinjiang region in the northwest of mainland China showed higher mass concentration (including the highest mass concentration) of pm2.5 from the year 2014 to 2019. There was a similar trend between the spatial variation of PM2.5 and it of AQI, PM10, CO, NO2, SO2 analyzed in the following content, but that of O3 would indicate a different trend compared with PM2.5. The highest annual mass concentration of PM2.5 was 161 μg·m-3, 201 μg·m-3, 204 μg·m-3, 187 μg·m-3, 217 μg·m-3, 183 μg·m-3 respectively in every separated year between 2014 and 2019. The highest mass concentration of PM2.5 in the year 2018 and 2014 were the highest and lowest among the six years, reaching 217 μg·m-3 and 161 μg·m-3 respectively. In addition, this index represented increase and then decrease, followed by an increase and decrease again, a kind of S-shaped change with time. The temporal variation of PM2.5 had a similar trend to that of AQI, PM10. On the other hand, from the perspective of the annual highest mass concentration of PM2.5, it can be analyzed that the atmospheric condition of mainland China had not been authentically improved until the year 2019.

**Figure 1.**
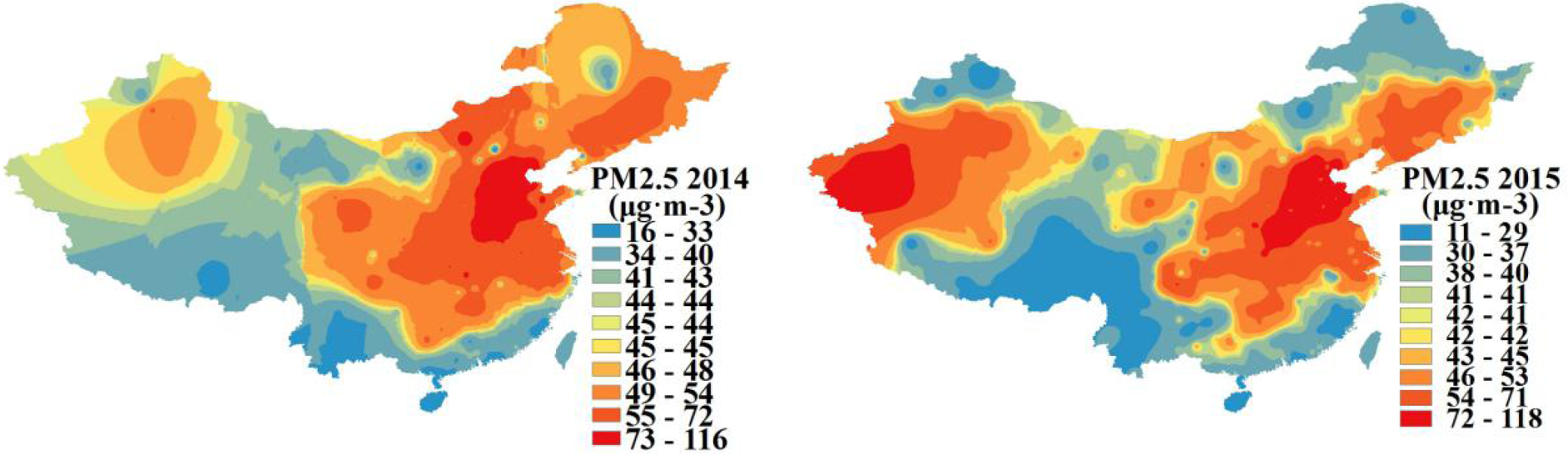
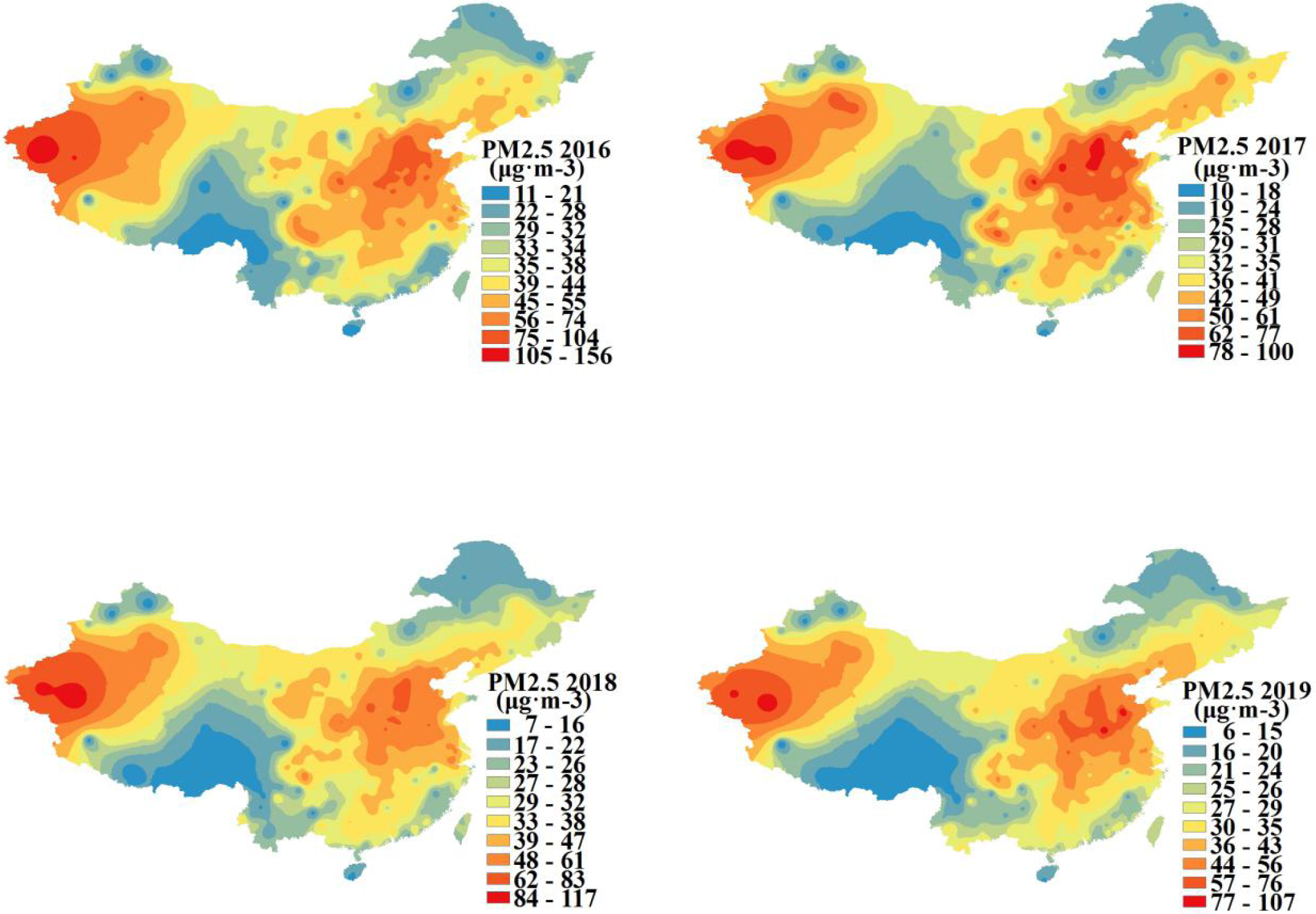
The spatial-temporal variation of the mass concentration of PM2.5 in mainland China from 2014 to 2019.

Table 1 indicates the ten cities with higher mass concentration of PM2.5 among 370 cities over the six-year period from 2014 to 2019, from which it can be seen that the proportions represented by some cities located in Beijing-Tianjin-Hebei region and Xinjiang region were about 5/6 and 1/6 respectively, with a lower proportion made up by some cities bordering Beijing-Tianjin-Hebei region. Among these records, Baoding and Xingtai, two cities situated in Beijing-Tianjin-Hebei region, combined with Kashi and Hetian, two areas situated in Xinjiang region, ranked the top of the list five to six times in the six years.

**Table 1.**
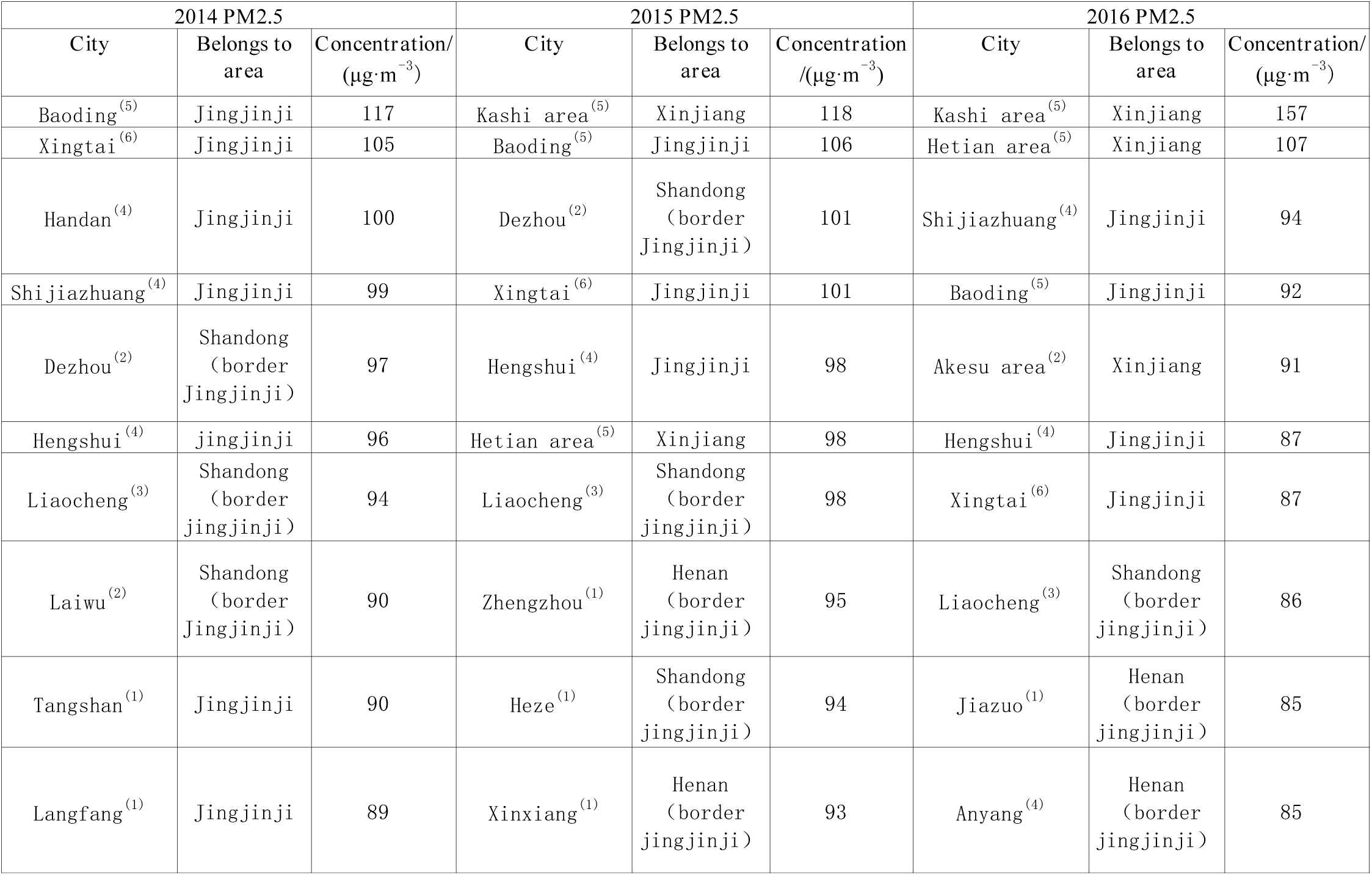
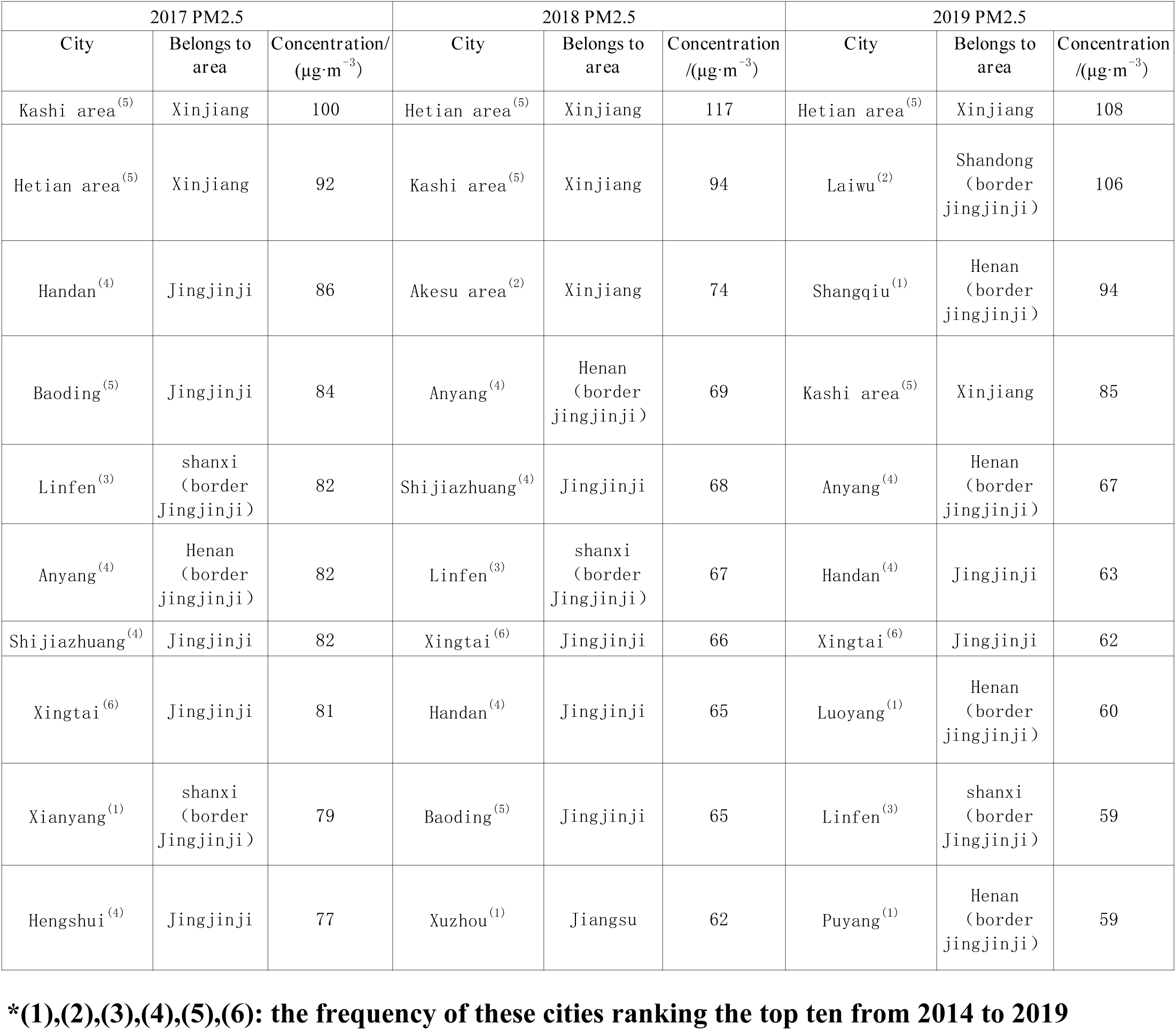
Ten cities with higher mass concentration of PM2.5 among 370 cities from 2014 to 2019.

From Figure 2, it can be seen that the Beijing-Tianjin-Hebei region and its surrounding region in the northeast of mainland China and Xinjiang region in the northwest of mainland China showed higher mass concentration (including the highest mass concentration) of AQI from the year 2014 to 2019. There was a similar trend between the spatial variation of AQI and it of the above PM2.5 and the following PM10, CO, NO2, SO2, but it of O3 would indicate a different trend compared with AQI. The highest mass concentration of AQI was 161, 201, 204,187, 217,183 respectively in every separated year between 2014 and 2019. The highest AQI in the year 2018 and 2014 were the highest and lowest among the six years, reaching 217 μg·m-3 and 161 μg·m-3 respectively. In addition, this index represented increase and then decrease, followed by an increase and decrease again, a kind of S-shaped change with time. The temporal variation of AQI had a similar trend to that of the above PM2.5 and the following PM10. On the other hand, from the perspective of the annual highest AQI, it can be analyzed that the atmospheric condition of mainland China had not been authentically improved until the year 2019.

**Figure 2.**
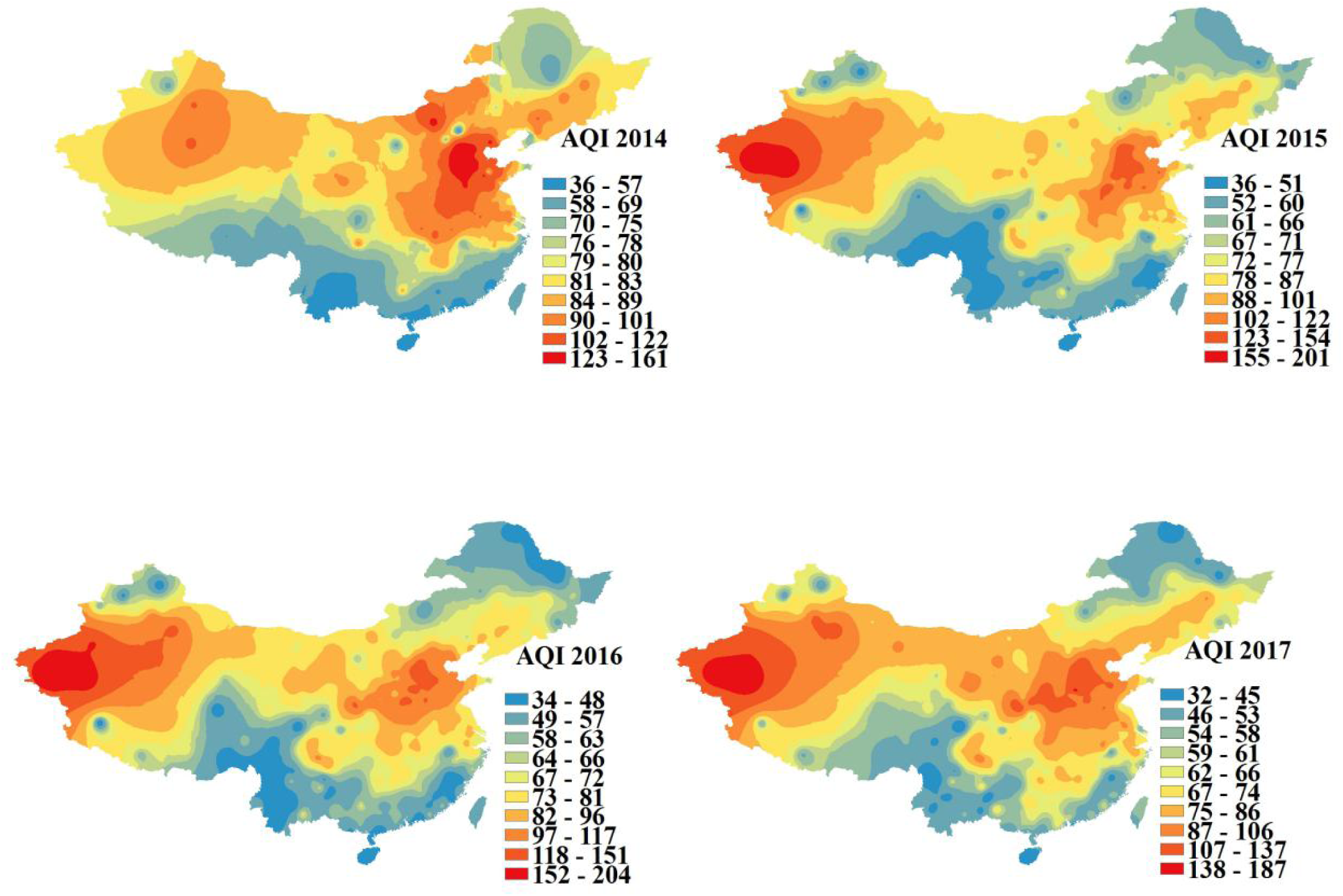
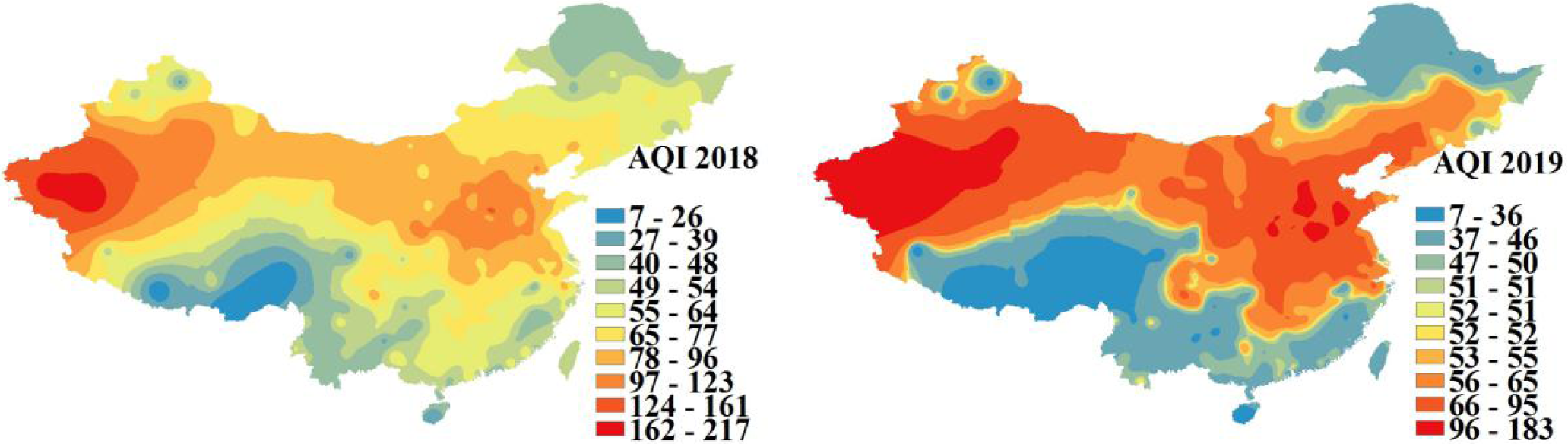
The spatial-temporal variation of the annual average AQI in mainland China from 2014 to 2019.

According to Table 2 which indicates the ten cities with higher mass concentration of AQI among 370 cities over the six-year period from 2014 to 2019, it can be seen that the proportions represented by some cities located in Beijing-Tianjin-Hebei region and Xinjiang region were almost the same, at about 1/2, with a lower proportion made up by some cities bordering Beijing-Tianjin-Hebei region. Among these records, Baoding, Xingtai and Handan, three cities situated in Beijing-Tianjin-Hebei region, combined with Kashi, Hetian, Aksu and Kezhou, four areas situated in Xinjiang region, ranked the top of the list four to five times in the six years.

**Table 2.** Ten cities with higher annual average AQI and their affiliated regions from 2014 to 2019.

| 2014 AQI |  |  | 2015 AQI |  |  | 2016 AQI |  |  |
| --- | --- | --- | --- | --- | --- | --- | --- | --- |
| City | Belongs to area | AQI | City | Belongs to area | AQI | City | Belongs to area | AQI |
| Baoding<sup>(4)</sup> | Jingjinji | 162 | Hetian area<sup>(5)</sup> | Xinjiang | 201 | Kashi area<sup>(5)</sup> | Xinjiang | 205 |
| Xingtai<sup>(5)</sup> | Jingjinji | 146 | Kashi area<sup>(5)</sup> | Xinjiang | 198 | Hetian area<sup>(5)</sup> | Xinjiang | 184 |
| Hengshui<sup>(3)</sup> | Jingjinji | 143 | Akesu area<sup>(5)</sup> | Xinjiang | 148 | Akesu area<sup>(5)</sup> | Xinjiang | 155 |
| Handan<sup>(4)</sup> | Jingjinji | 140 | Baoding<sup>(4)</sup> | Jingjinji | 144 | Kezhou area<sup>(5)</sup> | Xinjiang | 147 |
| Shijiazhuang<sup>(4)</sup> | Jingjinji | 138 | Hengshui<sup>(3)</sup> | Jingjinji | 141 | Kuerle<sup>(3)</sup> | Xinjiang | 133 |
| Dezhou<sup>(3)</sup> | Shandong<br>(border<br>Jingjinji) | 131 | Dezhou<sup>(3)</sup> | Shandong<br>(border<br>Jingjinji) | 139 | Shijiazhuang<sup>(3)</sup> | Jingjinji | 131 |
| Liaozheng<sup>(2)</sup> | Shandong<br>(border<br>Jingjinji) | 130 | Xingtai<sup>(5)</sup> | Jingjinji | 138 | Baoding<sup>(4)</sup> | Jingjinji | 128 |
| Langfang<sup>(1)</sup> | Jingjinji | 127 | Kezhou area<sup>(5)</sup> | Xinjiang | 137 | Tulufan area<sup>(4)</sup> | Xinjiang | 127 |
| Tangshan<sup>(1)</sup> | Jingjinji | 126 | Zhengzhou<sup>(1)</sup> | Henan<br>(border<br>jingjinji) | 135 | Hengshui<sup>(3)</sup> | Jingjinji | 126 |
| Heze<sup>(1)</sup> | Shandong<br>(border<br>Jingjinji) | 125 | Liaozheng<sup>(2)</sup> | Shandong<br>(border<br>Jingjinji) | 134 | Dezhou<sup>(3)</sup> | Shandong<br>(border<br>Jingjinji) | 123 |
| 2017 AQI |  |  | 2018 AQI |  |  | 2019 AQI |  |  |
| City | Belongs to area | AQI | City | Belongs to area | AQI | City | Belongs to area | AQI |
| Hetian area<sup>(5)</sup> | Xinjiang | 187 | Hetian area<sup>(5)</sup> | Xinjiang | 217 | Hetian area<sup>(5)</sup> | Xinjiang | 184 |
| Kashi area<sup>(5)</sup> | Xinjiang | 162 | Kashi area<sup>(5)</sup> | Xinjiang | 180 | Kashi area<sup>(5)</sup> | Xinjiang | 152 |
| Akesu area<sup>(5)</sup> | Xinjiang | 135 | Kezhou area<sup>(5)</sup> | Xinjiang | 155 | Laiwu<sup>(1)</sup> | Shandong<br>(border<br>Jingjinji) | 142 |
| Handan<sup>(4)</sup> | Jingjinji | 126 | Akesu area<sup>(5)</sup> | Xinjiang | 153 | Shangqiu<sup>(1)</sup> | Henan<br>(border<br>jingjinji) | 134 |
| Anyang<sup>(3)</sup> | Henan<br>(border<br>jingjinji) | 123 | Tulufan area<sup>(4)</sup> | Xinjiang | 122 | Akesu area<sup>(5)</sup> | Xinjiang | 120 |
| Xingtai<sup>(5)</sup> | Jingjinji | 122 | Kuerle<sup>(3)</sup> | Xinjiang | 121 | Kezhou area<sup>(5)</sup> | Xinjiang | 118 |
| Baoding<sup>(4)</sup> | Jingjinji | 122 | Anyang<sup>(3)</sup> | Henan<br>(border<br>jingjinji) | 115 | Kuerle<sup>(3)</sup> | Xinjiang | 116 |
| Shijiazhuang<sup>(3)</sup> | Jingjinji | 122 | Xingtai<sup>(5)</sup> | Jingjinji | 114 | Tulufan area<sup>(4)</sup> | Xinjiang | 112 |
| Xianyang<sup>(2)</sup> | shanxi<br>(border<br>Jingjinji) | 121 | Handan<sup>(4)</sup> | Jingjinji | 114 | Anyang<sup>(3)</sup> | Henan<br>(border<br>jingjinji) | 106 |
| Kezhou area<sup>(5)</sup> | Xinjiang | 120 | Wujiaqu<sup>(1)</sup> | Xinjiang | 113 | Handan<sup>(4)</sup> | Jingjinji | 102 |
| Tulufan area<sup>(4)</sup> | Xinjiang | 120 | Xianyang<sup>(2)</sup> | Shanxi<br>(border<br>Jingjinji) | 112 | Xingtai<sup>(5)</sup> | Jingjinji | 100 |

**\*(1),(2),(3),(4),(5),(6): the frequency of these cities ranking the top ten from 2014 to 2019**

From Figure 3, it can be seen that the Beijing-Tianjin-Hebei region and its surrounding region in the northeast of mainland China and Xinjiang region in the northwest of mainland China showed higher mass concentration (including the highest mass concentration) of PM10 from the year 2014 to 2019. There was a similar trend between the spatial variation of PM10 and it of the above PM2.5 and AQI and the following CO, NO2 and SO2, but it of the following O3 would indicate a different trend compared with PM10. The highest mass concentration of PM10 was 196 μg·m-3, 340 μg·m-3, 434 μg·m-3, 317 μg·m-3, 432 μg·m-3, 334 μg·m-3 respectively in every separated year between 2014 and 2019. The highest PM10 in the year 2016 and 2014 were the highest and lowest among the six years, reaching 434 μg·m-3 and 196 μg·m-3 respectively. In addition, this index represented increase and then decrease, followed by an increase and decrease again, a kind of S-shaped change with time. The temporal variation of PM10 had a similar trend to that of the above PM2.5 and AQI. On the other hand, from the perspective of the annual highest mass concentration of PM10, it can be analyzed that the atmospheric condition of mainland China had not been authentically improved until the year 2019.

**Figure 3.**
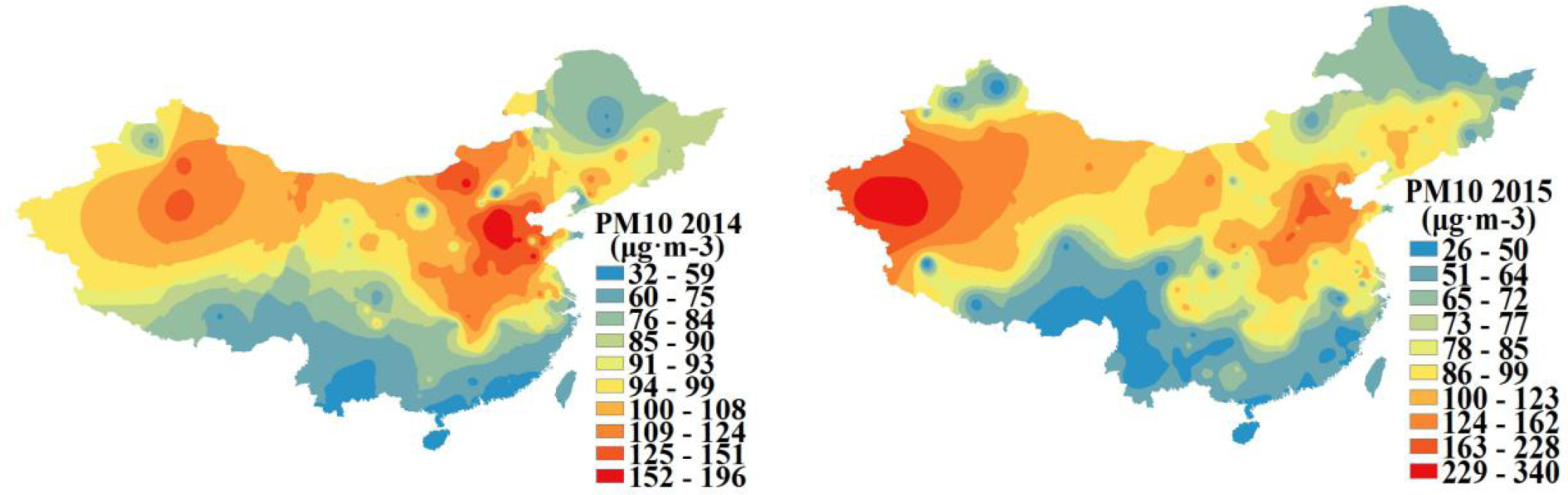
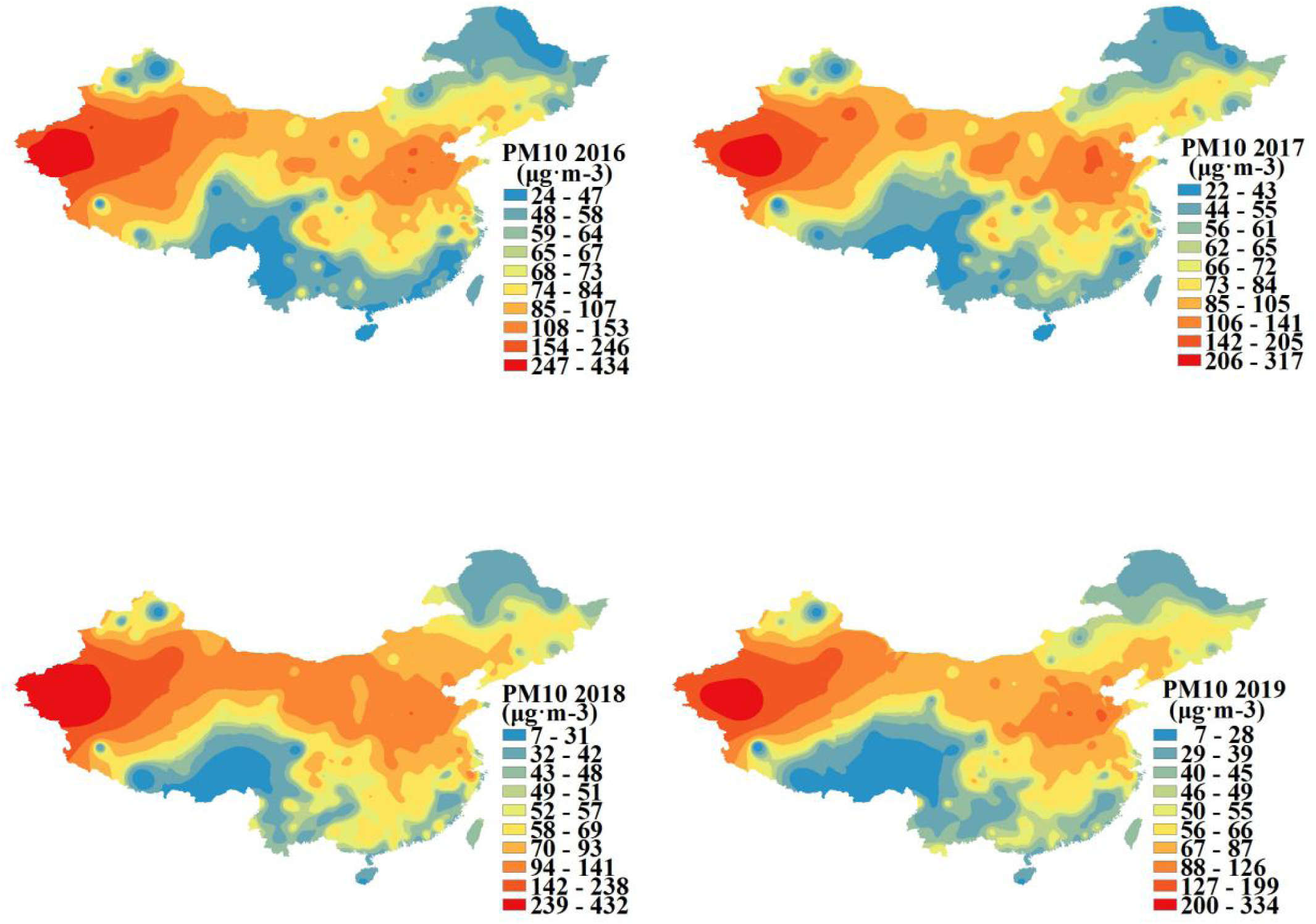
The spatial-temporal variation of the annual average mass concentration of PM10 in mainland China from 2014 to 2019.

According to Table 3 which indicates the ten cities with higher mass concentration of PM10 among 370 cities over the six-year period from 2014 to 2019, it can be seen that the proportions represented by Beijing-Tianjin-Hebei region and Xinjiang region were about 1/2, with a lower proportion made up by some areas bordering Beijing-Tianjin-Hebei region. Among these records, Handan, a city situated in Beijing-Tianjin-Hebei region, combined with Kashi, Hetian, Aksu and Kezhou, four areas situated in Xinjiang region, ranked the top of the list five to six times in the six years.

**Table 3.**
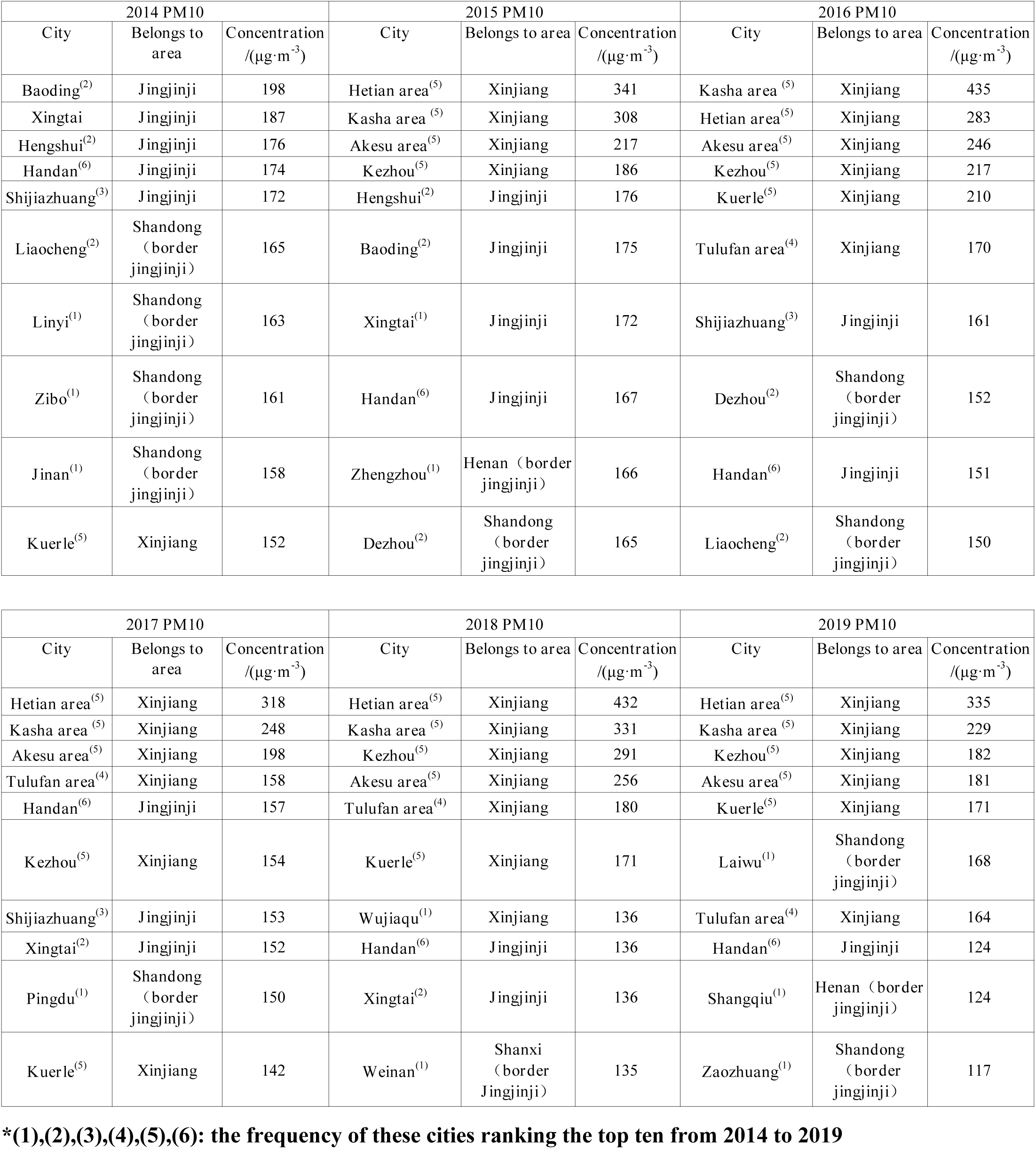
Ten cities with higher annual average mass concentration of PM10 and their affiliated regions from 2014 to 2019.

Similarly, it can be seen, from Figure 4, that the Beijing-Tianjin-Hebei region and its surrounding region in the northeast of mainland China and Xinjiang region in the northwest of mainland China showed higher mass concentration (including the highest mass concentration) of CO from the year 2014 to 2019. There was a similar trend between the spatial variation of CO and it of PM2.5, PM10, AQI, NO2 and SO2 acquired in the above and following contents, but it of the following O3 would indicate a different trend compared with CO. The highest mass concentration of CO was 2.2 μg·m-3, 2.5 μg·m-3, 2.4 μg·m-3,2 μg·m-3, 1.7 μg·m-3,1.1 μg·m-3 respectively in every separated year between 2014 and 2019. The highest mass concentration of CO in the year 2015 and 2019 were the highest (2.5 μg·m-3) and lowest (1.1 μg·m-3) among the six years. It also shows that this indicator had experienced a dramatic decline by the year 2019, which was less than the half of the figure in 2015. On the other hand, the temporal variation of CO witnessed a similar trend to that of the following NO2 and SO2. In addition, what is worth pointing is that extraordinarily high concentrations arose in Xinjiang region from 2018 to 2019. From the perspective of the annual highest mass concentration of CO, it can be analyzed that the atmospheric condition of mainland China had not been authentically improved until the year 2019.

**Figure 4.**
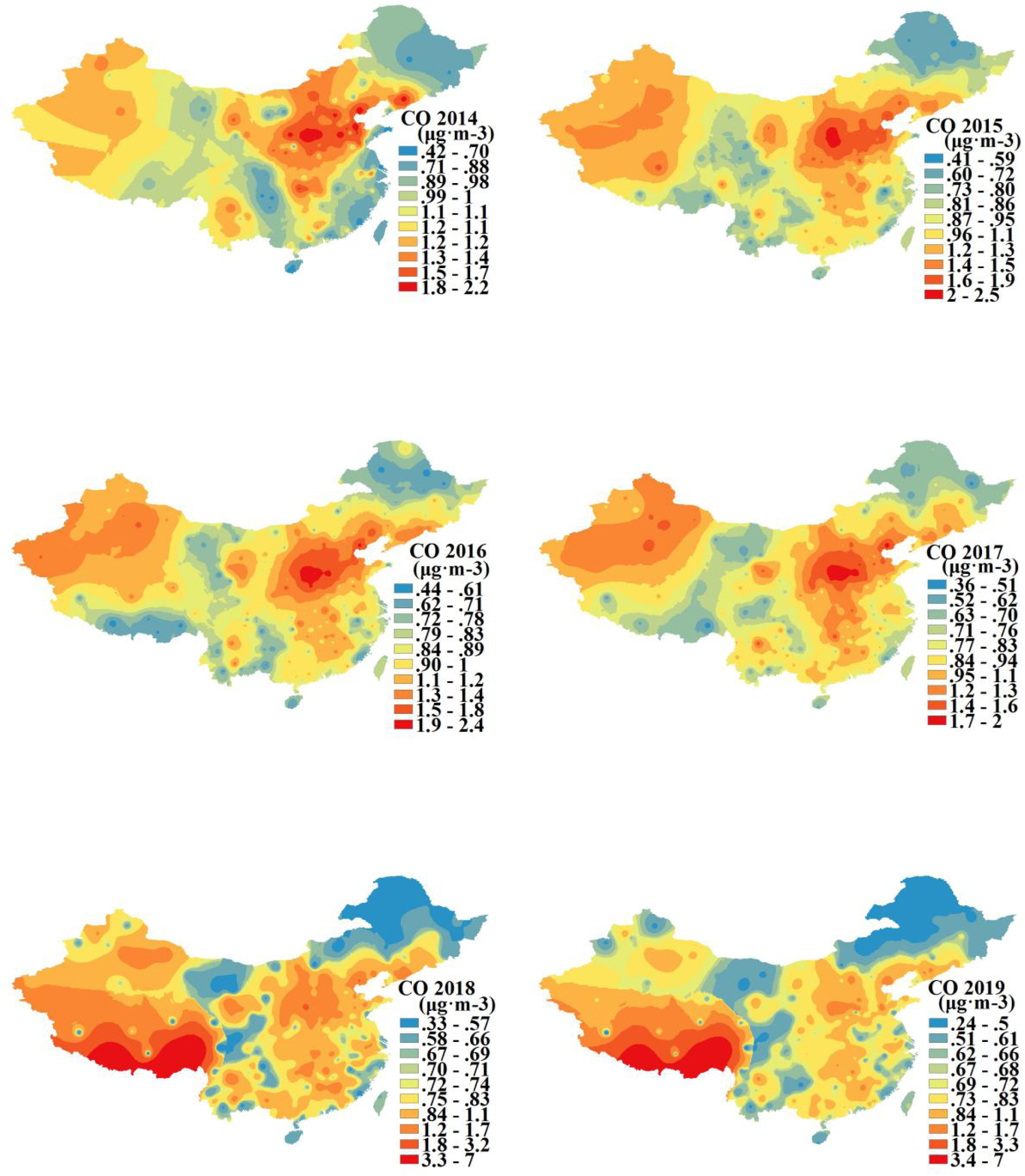
The spatial-temporal variation of the mass concentration of CO in mainland China from 2014 to 2019.

The top ten cities with higher annual mass concentration of CO from 2014 to 2019 and their affiliated regions are shown in Table 4. The results indicate that 1/3 of them were occupied by Shanxi, and Beijing-Tianjin-Hebei region and its neighboring areas like Shandong and Henan represented 1/2. To be more specific, Linfen and Changzhi, two cities situated in Shanxi province, combined with Anyang, a city situated in Henan province, ranked on the list of cities with top ten mass concentration of CO consecutively in the six years.

**Table 4.** Ten cities with higher annual average mass concentration of CO and their affiliated regions from 2014 to 2019.

| 2014 CO |  |  | 2015 CO |  |  | 2016 CO |  |  |
| --- | --- | --- | --- | --- | --- | --- | --- | --- |
| City | Belongs to area | Concentration $/(μg·m^{-3})$ | City | Belongs to area | Concentration $/(μg·m^{-3})$ | City | Belongs to area | Concentration $/(μg·m^{-3})$ |
| Penglai(2) | Shandong (border Jingjinji) | 2.25 | Lvliang(1) | Shanxi (border Jingjinji) | 2.75 | Linfen(6) | Shanxi (border Jingjinji) | 2.52 |
| Linfen(6) | Shanxi (border Jingjinji) | 2.20 | Linfen(6) | Shanxi (border Jingjinji) | 2.36 | Tangshan(6) | Jingjinji | 2.27 |
| Binzhou(2) | Shandong (border Jingjinji) | 2.13 | Tangshan(6) | Jingjinji | 2.13 | Anyang(5) | Henan (border jingjinji) | 2.17 |
| Zibo(4) | Shandong (border Jingjinji) | 2.11 | Zibo(4) | Shandong (border Jingjinji) | 2.11 | Yuncheng(3) | Shanxi (border Jingjinji) | 2.06 |
| Benxi(1) | Liaoning (border Jingjinji) | 2.09 | Binzhou(2) | Shandong (border Jingjinji) | 2.10 | Jincheng(4) | Shanxi (border Jingjinji) | 2.00 |
| Tangshan(6) | Jingjinji | 2.06 | Naqu area(1) | Xinjiang | 2.09 | Luoyang(1) | Henan (border jingjinji) | 2.00 |
| Changzhi(5) | Shanxi (border Jingjinji) | 2.05 | Anyang(5) | Henan (border jingjinji) | 2.06 | Changzhi(5) | Shanxi (border Jingjinji) | 2.00 |
| Liaocheng(1) | Shandong (border Jingjinji) | 2.00 | Zhoukou(1) | Henan (border jingjinji) | 2.02 | Hebi(2) | Henan (border jingjinji) | 1.96 |
| Yanan(1) | Shanxi (border Jingjinji) | 1.93 | Yuncheng(3) | Shanxi (border Jingjinji) | 1.96 | Zibo(4) | Shandong (border Jingjinji) | 1.90 |
| Baoding(1) | Jingjinji | 1.93 | Changzhi(5) | Shanxi (border Jingjinji) | 1.96 | Yilihasake(1) | Xinjiang | 1.85 |

| 2017 CO |  |  | 2018 CO |  |  | 2019 CO |  |  |
| --- | --- | --- | --- | --- | --- | --- | --- | --- |
| City | Belongs to area | Concentration $/(μg·m^{-3})$ | City | Belongs to area | Concentration $/(μg·m^{-3})$ | City | Belongs to area | Concentration $/(μg·m^{-3})$ |
| Linfen(6) | Shanxi (border Jingjinji) | 2.07 | Linfen(6) | Shanxi (border Jingjinji) | 1.76 | Laiwu(1) | Shandong (border Jingjinji) | 1.53 |
| Jincheng(4) | Shanxi (border Jingjinji) | 1.98 | Tangshan(6) | Jingjinji | 1.69 | Linfen(6) | Shanxi (border Jingjinji) | 1.46 |
| Tangshan(6) | Jingjinji | 1.96 | Panzhuhua(2) | Sichuan | 1.53 | Tangshan(6) | Jingjinji | 1.37 |
| Anyang(5) | Henan (border jingjinji) | 1.85 | Anyang(5) | Henan (border jingjinji) | 1.53 | Datong(2) | Shanxi (border Jingjinji) | 1.35 |
| Dekehasake(2) | Xinjiang | 1.83 | Dekehasake(2) | Xinjiang | 1.46 | Changzhi(5) | Shanxi (border Jingjinji) | 1.25 |
| Zibo(4) | Shandong (border Jingjinji) | 1.76 | Loudi(1) | Hunan | 1.45 | Ganzhou(2) | Jiangxi | 1.24 |
| Yuncheng(3) | Shanxi (border Jingjinji) | 1.71 | Datong(2) | Shanxi (border Jingjinji) | 1.42 | Jincheng(4) | Shanxi (border Jingjinji) | 1.23 |
| Hebi(2) | Henan (border jingjinji) | 1.68 | Ganzhou(2) | Jiangxi | 1.42 | Anyang(5) | Henan (border jingjinji) | 1.21 |
| Changzhi(5) | Shanxi (border Jingjinji) | 1.67 | Xining(1) | Qinghai | 1.41 | Panzhuhua(2) | Sichuan | 1.20 |
| Xingtai | Jingjinji | 1.67 | Jincheng(4) | Shanxi (border Jingjinji) | 1.38 | Penglai(2) | Shandong (border Jingjinji) | 1.19 |

\*(1),(2),(3),(4),(5),(6): the frequency of these cities ranking the top ten from 2014 to 2019

From Figure 5, it can be seen that northern cities or areas with higher concentration (including the highest mass concentration) of NO2 are also mostly situated in the Beijing-Tianjin-Hebei region and Xinjiang region from the year 2014 to 2017. Simultaneously, the concentration of NO2 took a high level position in Yangtze River Delta region, Pearl River Delta region and Sichuan Basin. It should be pointed that the distribution of NO2 was more extensive than other parameters, especially in the region where local economy was more developed such as Yangtze River Delta region, Pearl River Delta region and Sichuan Basin. There was a similar trend between the spatial variation of NO2 and it of the above PM2.5, PM10, AQI, CO and the following SO2, but it of the following O3 would indicate an opposite trend compared with CO. The highest mass concentration of NO2 was 56 μg·m-3, 59 μg·m-3, 60 μg·m-3,58 μg·m-3, 54 μg·m-3,50 μg·m-3 respectively in every separated year between 2014 and 2019. The highest mass concentration of NO2 in the year 2016 and 2019 were the highest (60 μg·m-3) and lowest (50 μg·m-3) among the six years. On the other hand, the temporal variation of NO2 witnessed a similar trend to that of the above CO and the following SO2, but a different trend to that of the following O3. From the perspective of the annual highest mass concentration of NO2, it can be analyzed that the atmospheric condition of mainland China had not been authentically improved until the year 2019.

**Figure 5.**
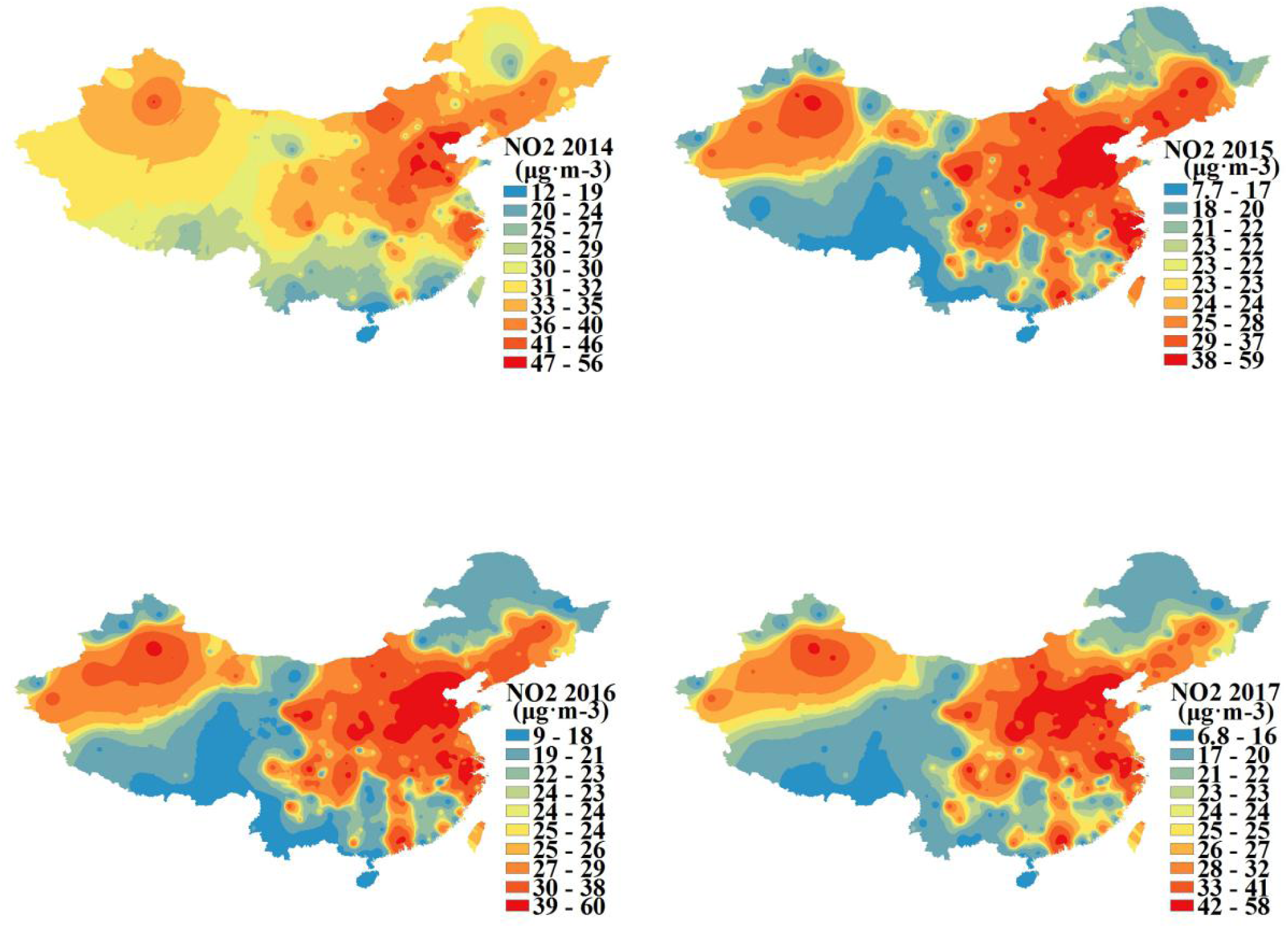
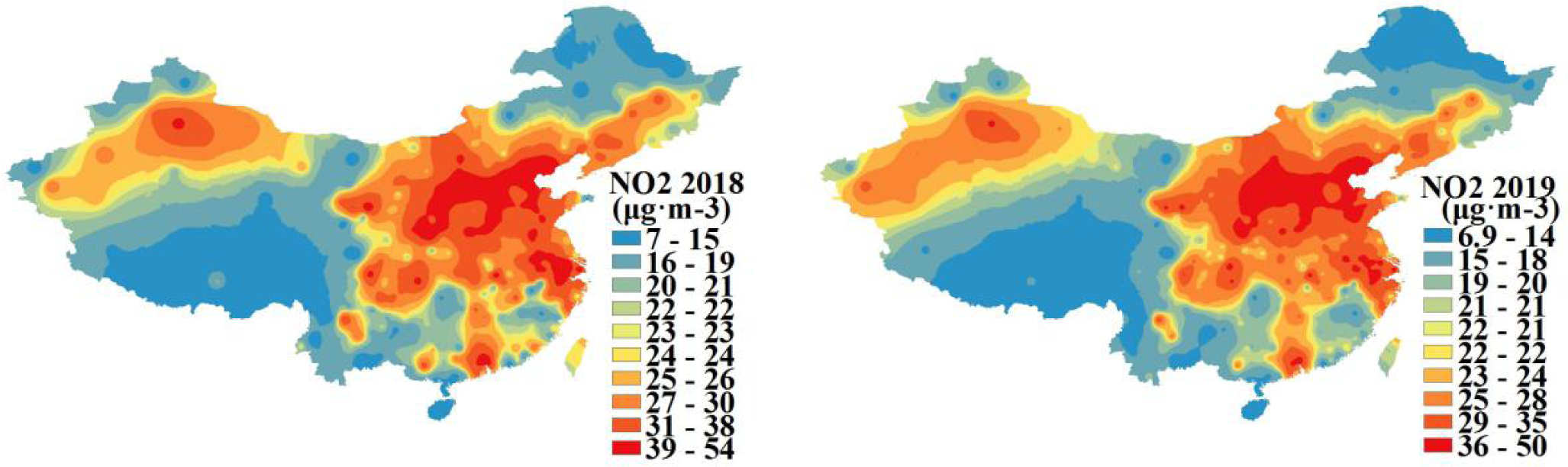
The spatial-temporal variation of the mass concentration of NO2 in mainland China from 2014 to 2019.

Table 5 indicates the cities whose annual average mass concentration of NO2 ranked the top then from 2014 to 2019 and their affiliated regions.It can be seen that the number of these cities in Beijing-Tianjin-Hebei region and its neighboring areas accounted more than 2/3.Tangshan and Xingtai, two cities bordering.Beijing-Tianjin-Hebei region, ranked on the list of cities with top ten mass concentration of NO2 consecutively in the six years. Also, Zhengzhou, the provincial capital of Henan province bordering Beijing-Tianjin-Hebei region, similarly ranked on this list up to five times in these years.

**Table 5.** Ten cities with higher annual average mass concentration of NO2 and their affiliated regions from 2014 to 2019.

their affiliated regions from 2014 to 2019
| 2014 NO2 |  |  | 2015 NO2 |  |  | 2016 NO2 |  |  |
| --- | --- | --- | --- | --- | --- | --- | --- | --- |
| City | Belongs to area | Concentration<br>/($\mu\text{g}\cdot\text{m}^{-3}$) | City | Belongs to area | Concentration<br>/($\mu\text{g}\cdot\text{m}^{-3}$) | City | Belongs to area | Concentration<br>/($\mu\text{g}\cdot\text{m}^{-3}$) |
| Zibo(3) | Shandong<br>(border<br>jingjinji) | 65 | Zibo(3) | Shandong<br>(border<br>jingjinji) | 61 | Xingtai(6) | Jingjinji | 61 |
| Tangshan(6) | Jingjinji | 58 | Tangshan(6) | Jingjinji | 60 | Tangshan(6) | Jingjinji | 58 |
| Linyi(1) | Shandong<br>(border<br>jingjinji) | 57 | Xingtai(6) | Jingjinji | 59 | Baoding(3) | Jingjinji | 57 |
| Xingtai(6) | Jingjinji | 56 | Zhengzhou(5) | Henan<br>(border<br>jingjinji) | 55 | Handan(2) | Jingjinji | 54 |
| Jinan(3) | Shandong<br>(border<br>jingjinji) | 53 | Baoding(3) | Jingjinji | 53 | Huzhou(3) | Jiangsu | 54 |
| Baoding(3) | Jingjinji | 52 | Linxiashou(1) | Gansu | 53 | Shijiazhuang(2) | Jingjinji | 54 |
| Beijing(1) | Jingjinji | 52 | Suzhou(2) | Jiangsu | 51 | Zhengzhou(5) | Henan<br>(border<br>jingjinji) | 53 |
| Suzhou(2) | Jiangsu | 51 | Jinan(3) | Shandong<br>(border<br>jingjinji) | 51 | Zibo(3) | Shandong<br>(border<br>jingjinji) | 52 |
| Laiwu(2) | Shandong<br>(border<br>jingjinji) | 51 | Zhuji(1) | Zhejiang | 51 | Hebi(1) | Henan<br>(border<br>jingjinji) | 52 |
| Qinhuangdao(2) | Shandong<br>(border<br>jingjinji) | 50 | Xinxiang(2) | Henan<br>(border<br>jingjinji) | 51 | Wulumuqi(1) | Xinjiang | 52 |

| 2017 NO2 |  |  | 2018 NO2 |  |  | 2019 NO2 |  |  |
| --- | --- | --- | --- | --- | --- | --- | --- | --- |
| City | Belongs to area | Concentration /($\mu\text{g}\cdot\text{m}^{-3}$) | City | Belongs to area | Concentration /($\mu\text{g}\cdot\text{m}^{-3}$) | City | Belongs to area | Concentration /($\mu\text{g}\cdot\text{m}^{-3}$) |
| Tangshan(6) | Jingjinji | 58 | Huzhou(3) | Jiangsu | 54 | Laiwu(2) | Shandong (border jingjinji) | 51 |
| Xian(3) | Shan Xi | 58 | Tangshan(6) | Jingjinji | 54 | Tangshan(6) | Jingjinji | 49 |
| Xingtai(6) | Jingjinji | 56 | Xian(3) | Shan Xi | 53 | Yunlin(1) | Henan (border jingjinji) | 47 |
| Huzhou(3) | Jiangsu | 55 | Changzhou(1) | Jiangsu | 49 | Xian(3) | Shan Xi | 46 |
| Xianyang(2) | Shan Xi | 53 | Tianjin(2) | Jingjinji | 49 | Taiyuan(2) | Shanxi (border jingjinji) | 44 |
| Tianjin(2) | Jingjinji | 53 | Xingtai(6) | Jingjinji | 47 | Lvliang | Shanxi (border jingjinji) | 44 |
| Zhengzhou(5) | Henan (border jingjinji) | 52 | Xinxiang(2) | Henan (border jingjinji) | 47 | Huzhou(2) | Jiangsu | 44 |
| Weinan(1) | Shan Xi | 51 | Xianyang(2) | Shan Xi | 47 | Jinan(3) | Shandong (border jingjinji) | 43 |
| Handan(2) | Jingjinji | 51 | Taiyuan(2) | Shanxi (border jingjinji) | 46 | Xingtai(6) | Jingjinji | 42 |
| Shijiazhuang(2) | Jingjinji | 50 | Zhengzhou(5) | Henan (border jingjinji) | 46 | Zhengzhou(5) | Henan (border jingjinji) | 42 |

\*(1),(2),(3),(4),(5),(6): the frequency of these cities ranking the top ten from 2014 to 2019

From Figure 6, it can be seen that higher concentration (including the highest mass concentration) of O3 was mainly distributed in Qinghai, Inner Mongolia in the central north of mainland China and Shandong province bordering Beijing-Tianjin-Hebei region on the right side from the year 2014 to 2019. In fact, the spatial variation of O3 experienced a different trend compared with other kinds of atmospheric pollution parameters on account that only O3 showed a high concentration in both Qinghai and Inner Mongolia regions. The highest mass concentration of O3 was 47 μg·m-3, 93 μg·m-3, 95 μg·m-3,100 μg·m-3, 100 μg·m-3,100 μg·m-3 respectively in every separated year between 2014 and 2019. The highest mass concentration of O3 in the year 2014 were the lowest (47 μg·m-3) among the six years and after 2015, the highest mass concentration of O3 was slightly higher in each year. On the other hand, the temporal variation of O3 had different trends compared with the above CO and NO2 and the following SO2. From the perspective of the annual highest mass concentration of O3, it can be analyzed that the atmospheric condition of mainland China had become worse by the year 2019.

**Figure 6.**
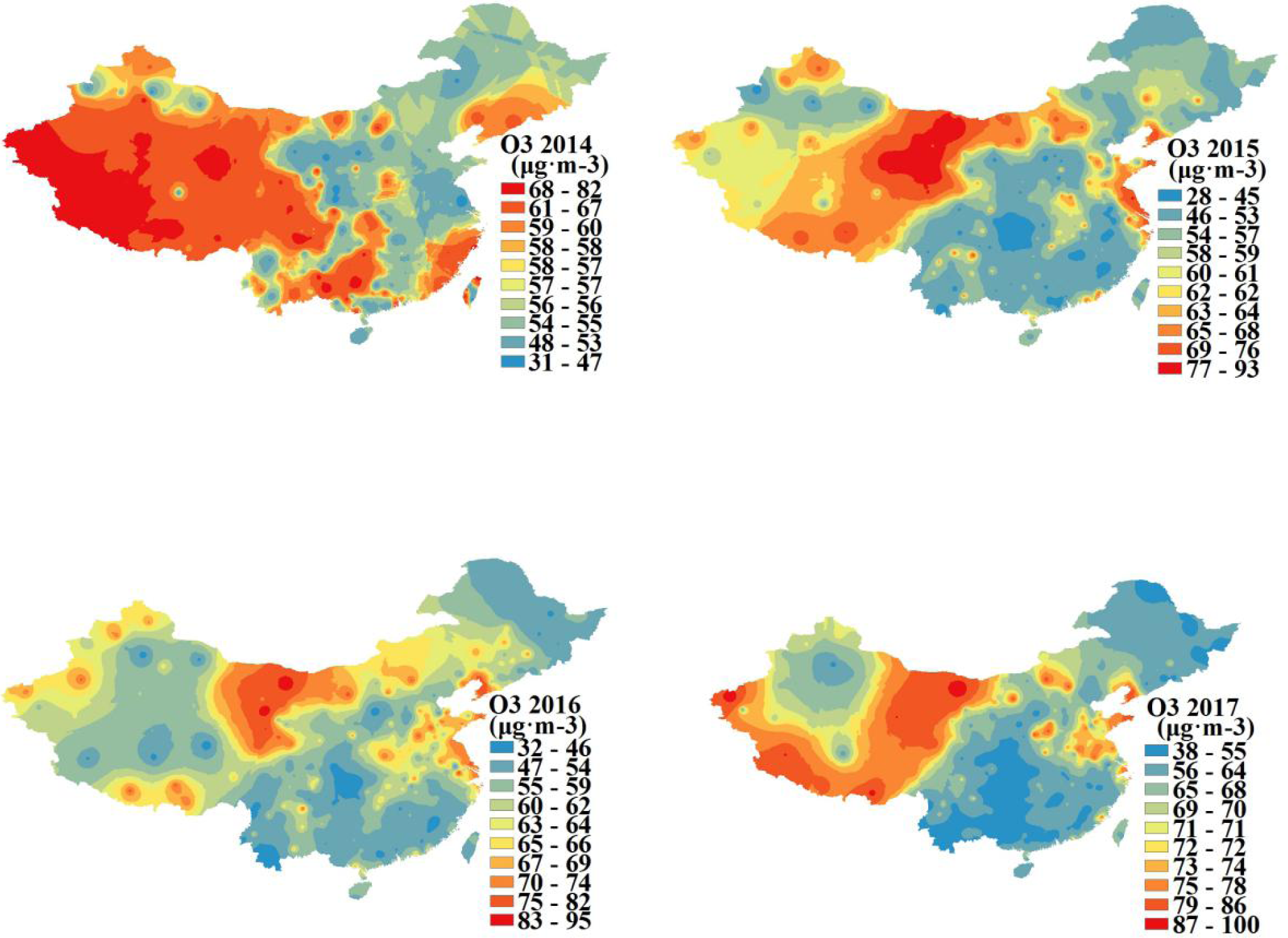
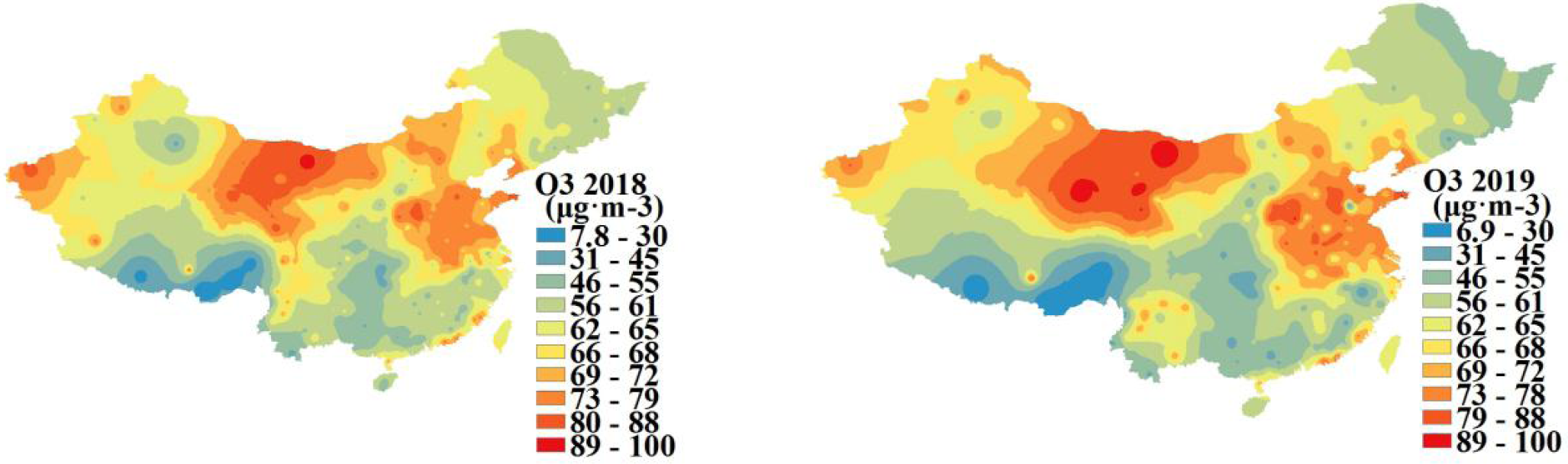
The spatial-temporal variation of the mass concentration of O3 in mainland China from 2014 to 2019.

Table 6 lists the cities with top ten annual mass concentration of O3 and their affiliated regions from the year 2014 to 2019, which indicates that more than 1/3 of them were situated in Qinghai and Inner Mongolia regions and 1/6 of them were situated in Shandong province bordering Beijing-Tianjin-Hebei region, with the rest distributed the neighboring regions like Gansu, Liaoning provinces, etc. Especially, in four out of the total six years, Haibeizhou in Qinghai and Alxa in Inner Mongolia were selected into the cities whose annual mass concentration was top ten.

**Table 6.** Ten cities with higher annual average mass concentration of O3 and their affiliated regions from 2014 to 2019.

| 2014 O<sub>3</sub> |  |  | 2015 O<sub>3</sub> |  |  | 2016 O<sub>3</sub> |  |  |
| --- | --- | --- | --- | --- | --- | --- | --- | --- |
| City | Belongs to area | Concentration/<br>($\mu\text{g}\cdot\text{m}^{-3}$) | City | Belongs to area | Concentration/<br>($\mu\text{g}\cdot\text{m}^{-3}$) | City | Belongs to area | Concentration/<br>($\mu\text{g}\cdot\text{m}^{-3}$) |
| Rongcheng(3) | Shandong<br>(border<br>jingjinji) | 93 | Haixizhou(3) | Qinghai | 97 | Haibeizhou(4) | Qinghai | 95 |
| Weihai(4) | Shandong<br>(border<br>jingjinji) | 92 | Alashanmeng(4) | Neimenggu | 94 | Alashanmeng(4) | Neimenggu | 95 |
| Penglai(1) | Shandong<br>(border<br>jingjinji) | 87 | Hainanzhou(3) | Qinghai | 94 | Hainanzhou(3) | Qinghai | 83 |
| Linan(1) | Zhejiang | 85 | Haibeizhou(4) | Qinghai | 90 | Yixing(1) | Jiangsu | 82 |
| Jiaonan(2) | Shandong<br>(border<br>jingjinji) | 82 | Yanzheng(1) | Jiangsu | 82 | Guoluo Zhou(3) | Qinghai | 81 |
| Weifang(1) | Shandong<br>(border<br>jingjinji) | 81 | Jiayuguan(2) | Gansu | 81 | Yingkou(2) | Liaoning | 81 |
| Wendeng(1) | Shandong<br>(border<br>jingjinji) | 81 | Guoluo Zhou(3) | Qinghai | 81 | Jiayuguan(2) | Gansu | 81 |
| Dongying(1) | Shandong<br>(border<br>jingjinji) | 80 | Yingkou(2) | Liaoning | 80 | Eerduosi(2) | Neimenggu | 80 |
| Eerduosi(2) | Neimenggu | 80 | Lasa(1) | Xizang | 80 | Jiaonan(1) | Jiangsu | 80 |
| Qingdao(1) | Shandong<br>(border<br>jingjinji) | 80 | Weihai(4) | Shandong<br>(border<br>jingjinji) | 79 | Dalian(1) | Liaoning | 79 |

| 2017 NO2 |  |  | 2018 NO2 |  |  | 2019 NO2 |  |  |
| --- | --- | --- | --- | --- | --- | --- | --- | --- |
| City | Belongs to area | Concentration/<br>($\mu\text{g}\cdot\text{m}^{-3}$) | City | Belongs to area | Concentration/<br>($\mu\text{g}\cdot\text{m}^{-3}$) | City | Belongs to area | Concentration/<br>($\mu\text{g}\cdot\text{m}^{-3}$) |
| Enshizhou(1) | Hubei | 37 | Alashanmeng(4) | Neimenggu | 105 | Haixizhou(3) | Qinghai | 104 |
| Xiangxi | Hunan | 38 | Guoluozhou(3) | Qinghai | 90 | Alashanmeng(4) | Neimenggu | 102 |
| Qiandongnanzhou(1) | Guizhou | 38 | Kezhou(1) | Xinjiang | 89 | Haibeizhou(4) | Qinghai | 96 |
| Xishuangbanna(1) | Yunnan | 40 | Weihai(4) | Shandong<br>(border<br>jingjinji) | 89 | Weihai(4) | Shandong<br>(border<br>jingjinji) | 92 |
| Qianxinanzhou(1) | Guizhou | 40 | Haixizhou(3) | Qinghai | 88 | Hainanzhou(3) | Qinghai | 89 |
| Nujiang(1) | Yunnan | 41 | Jiaozuo(2) | Henan<br>(border<br>jingjinji) | 87 | Jiaozuo(2) | Henan<br>(border<br>jingjinji) | 89 |
| Chongqing(1) | Chongqing | 42 | Haibeizhou(4) | Qinghai | 87 | Rongcheng(3) | Shandong<br>(border<br>jingjinji) | 89 |
| Wuzhou(1) | Guangxi | 42 | Rongcheng(3) | Shandong<br>(border<br>jingjinji) | 85 | Jinchang(1) | Gansu | 86 |
| Hechi(1) | Jiangsu | 43 | Qingyang(1) | Gansu | 85 | Changzhi(2) | Shanxi<br>(border<br>jingjinji) | 85 |
| Pingxiang(1) | Jiangxi | 43 | Changzhi(2) | Shanxi<br>(border<br>jingjinji) | 84 | Jincheng(1) | Shanxi<br>(border<br>jingjinji) | 84 |

\*(1),(2),(3),(4),(5),(6): the frequency of these cities ranking the top ten from 2014 to 2019

Based on the combination of Figure 6 and Table 6, it can be seen that the distribution of O3 in mainland China was different from that of other atmospheric pollution parameters discussed in this paper. In fact, the concentration of O3 distributed in Qinghai and Inner Mongolia in North China was higher.

From Figure 7, it can be seen that the area with higher concentration (including the highest mass concentration) of SO2 was mainly situated in Shanxi and Shandong provinces nearby Beijing-Tianjin-Hebei region and a small part of areas bordering such places (e.g. Liaoning province and Inner Mongolia region) from the year 2014 to 2019. There was a similar trend between the spatial variation of SO2 and that of the above PM2.5, PM10, AQI, CO and NO2, but a different trend between the spatial variation of SO2 and that of the above O3. The highest mass concentration of SO2 was 83 μg·m-3, 76 μg·m-3, 87 μg·m-3,82 μg·m-3,48 μg·m-3,37 μg·m-3 in each separated year from 2014 to 2019. Among these, the highest concentration of SO2 was 87 μg·m-3 in 2016, which was higher than the other five years and more than two times as high as that in 2019, the lowest one with figure 37 μg·m-3. After the year 2018, the highest mass concentration of SO2 in atmosphere experienced a significant decline. On the other hand, the temporal variations of SO2 and the above CO and NO2 indicated similar trend but that of O3 was different. From the perspective of the annual highest mass concentration of SO2, it can be analyzed that the atmospheric condition of mainland China had been remarkably improved by the year 2019.

**Figure 7.**
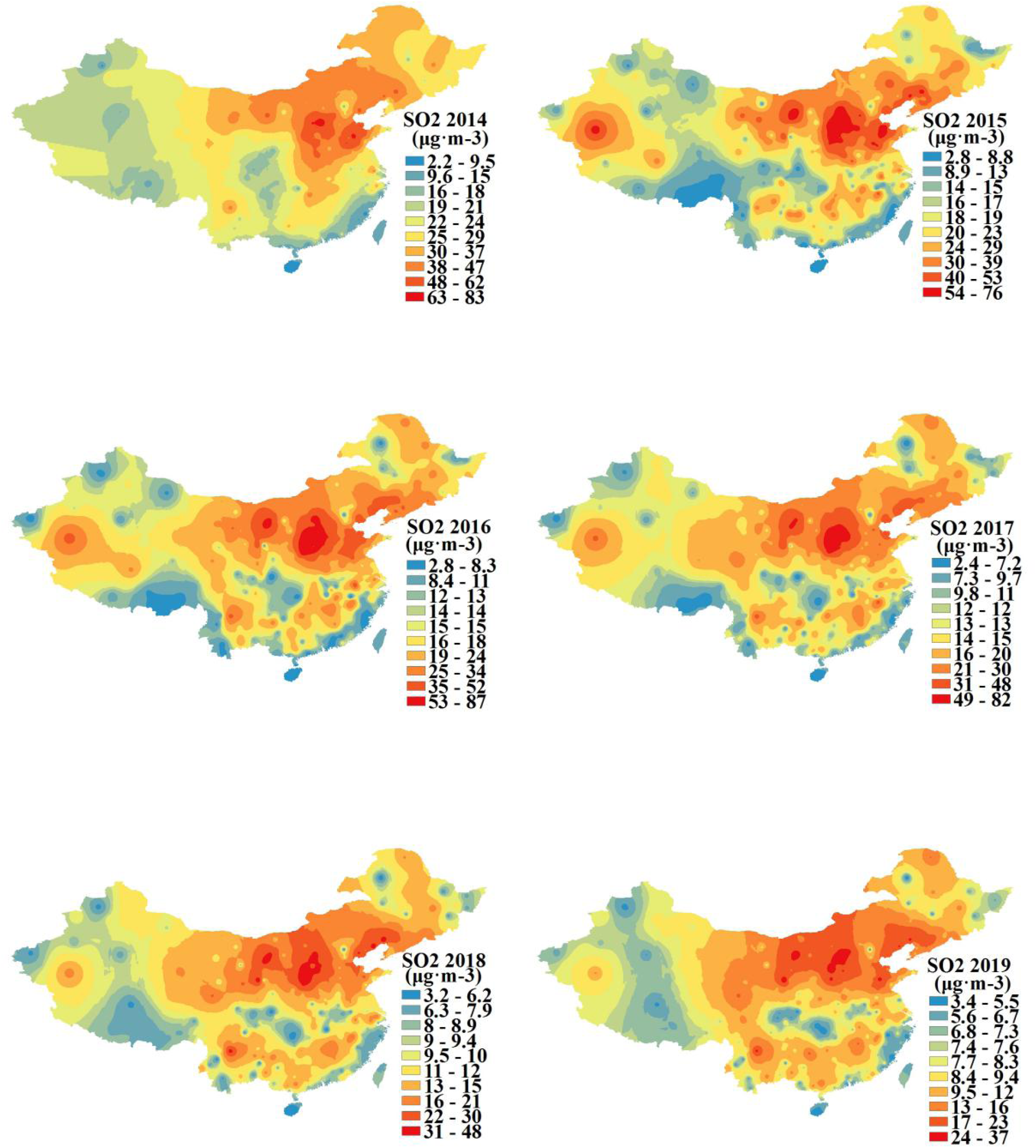
The spatial-temporal variation of the mass concentration of SO2 in mainland China from 2014 to 2019.

There are some cities with top ten annual mass concentration of SO2 from the year 2014 to 2019 and their affiliated regions in Table 7, which indicates that more than 1/2 of these areas existed in Shanxi province on the left side of Beijing-Tianjin-Hebei region, with 1/6 in Shandong province on the right side of Beijing-Tianjin-Hebei region, 1/6 in Ningxia and the rest in the areas bordering the above places, like Inner Mongolia and Liaoning Province. In six consecutive years, Linfen, located in Shanxi province, ranked the top ten cities with higher annual mass concentration of SO2, followed by Shijiazui in Ningxia, which was selected onto the list in five years.

**Table 3.** Ten cities with higher annual average mass concentration of SO2 and their affiliated regions from 2014 to 2019.

| 2014 SO2 |  |  | 2015 SO2 |  |  | 2016 SO2 |  |  |
| --- | --- | --- | --- | --- | --- | --- | --- | --- |
| City | Belongs to area | Concentration<br>/($\mu\text{g}\cdot\text{m}^{-3}$) | City | Belongs to area | Concentration<br>/($\mu\text{g}\cdot\text{m}^{-3}$) | City | Belongs to area | Concentration<br>/($\mu\text{g}\cdot\text{m}^{-3}$) |
| Zibo(2) | Shandong<br>(border<br>jingjinji) | 108 | Zibo(2) | Shandong<br>(border<br>jingjinji) | 82 | Jinzhong(4) | Shanxi<br>(border<br>jingjinji) | 87 |
| Yangquan(3) | Shanxi<br>(border<br>jingjinji) | 85 | Shuozhou(3) | Shanxi<br>(border<br>jingjinji) | 76 | Linfen(6) | Shanxi<br>(border<br>jingjinji) | 82 |
| Laiwu(2) | Shandong<br>(border<br>jingjinji) | 81 | Hetiandiqu(1) | Xinjiang | 72 | Shuozhou(3) | Shanxi<br>(border<br>jingjinji) | 67 |
| Taiyuan(4) | Shanxi<br>(border<br>jingjinji) | 66 | Jinzhong(4) | Shanxi<br>(border<br>jingjinji) | 72 | Shijiazui(5) | Ningxia | 67 |
| Jining | Shandong<br>(border<br>jingjinji) | 64 | Shijiazui(5) | Ningxia | 71 | Jincheng(1) | Shanxi<br>(border<br>jingjinji) | 66 |
| Shandong | Shandong<br>(border<br>jingjinji) | 64 | Taiyuan(4) | Shanxi<br>(border<br>jingjinji) | 68 | Yuncheng(2) | Shanxi<br>(border<br>jingjinji) | 66 |
| Zaozhuang | Shandong<br>(border<br>jingjinji) | 63 | Lvliang(5) | Shanxi<br>(border<br>jingjinji) | 68 | Taiyuan(4) | Shanxi<br>(border<br>jingjinji) | 65 |
| Linfen(6) | Shanxi<br>(border<br>jingjinji) | 63 | Zhangqiu(1) | Shandong<br>(border<br>jingjinji) | 64 | Lvliang(5) | Shanxi<br>(border<br>jingjinji) | 62 |
| Shijiazui(5) | Ningxia | 59 | Linfen(6) | Shanxi<br>(border<br>jingjinji) | 63 | Yangquan(3) | Shanxi<br>(border<br>jingjinji) | 61 |
| Shenyang(1) | Liaoning<br>(border<br>jingjinji) | 57 | Wuhai(4) | Neimenggu | 62 | Changzhi(1) | Shanxi<br>(border<br>jingjinji) | 61 |

| 2017 SO2 |  |  | 2018 SO2 |  |  | 2019 SO2 |  |  |
| --- | --- | --- | --- | --- | --- | --- | --- | --- |
| City | Belongs to area | Concentration<br>/($\mu\text{g}\cdot\text{m}^{-3}$) | City | Belongs to area | Concentration<br>/($\mu\text{g}\cdot\text{m}^{-3}$) | City | Belongs to area | Concentration<br>/($\mu\text{g}\cdot\text{m}^{-3}$) |
| Jinzhong(4) | Shanxi<br>(border<br>jingjinji) | 83 | Linfen(6) | Shanxi<br>(border<br>jingjinji) | 48 | Baiyin(2) | Ningxia | 39 |
| Linfen(6) | Shanxi<br>(border<br>jingjinji) | 79 | Baiyin(2) | Ningxia | 42 | Wuhai(4) | Neimenggu | 33 |
| Lvliang(5) | Shanxi<br>(border<br>jingjinji) | 68 | Lvliang(5) | Shanxi<br>(border<br>jingjinji) | 39 | Laiwu(2) | Shandong<br>(border<br>jingjinji) | 32 |
| Shijiazui(5) | Ningxia | 51 | Panzhihua(2) | Sichuan | 38 | Shuozhou(3) | Shanxi<br>(border<br>jingjinji) | 29 |
| Taiyuan(4) | Shanxi<br>(border<br>jingjinji) | 51 | Jinzhou(2) | Liaoning<br>(border<br>jingjinji) | 37 | Panzhihua(2) | Sichuan | 29 |
| Yinchuang(2) | Ningxia | 51 | Shijiazui(5) | Ningxia | 37 | Xinzhou(2) | Shanxi<br>(border<br>jingjinji) | 29 |
| Yuncheng(2) | Shanxi<br>(border<br>jingjinji) | 51 | Jinzhong(4) | Shanxi<br>(border<br>jingjinji) | 36 | Lvliang(5) | Shanxi<br>(border<br>jingjinji) | 29 |
| Wuhai(4) | Neimenggu | 51 | Yinchuang(2) | Ningxia | 36 | Datong(1) | Shanxi<br>(border<br>jingjinji) | 28 |
| Yangquan(3) | Shanxi<br>(border<br>jingjinji) | 49 | Huludao(1) | Liaoning<br>(border<br>jingjinji) | 36 | Linfen(6) | Shanxi<br>(border<br>jingjinji) | 27 |
| Xinzhou(2) | Shanxi<br>(border<br>jingjinji) | 48 | Wuhai(4) | Neimenggu | 34 | Jinzhou(2) | Liaoning<br>(border<br>jingjinji) | 27 |

\*(1),(2),(3),(4),(5),(6): the frequency of these cities ranking the top ten from 2014 to 2019

## 3 Conclusion

The spatial-temporal distribution law of atmospheric pollution in China is of persistence, concentricity and correlation between different atmospheric pollution parameters. From the year 2014 to 2019, such atmospheric pollution parameters as PM2.5, AQI, PM10, CO, NO2 and SO2 were mainly concentrated in Beijing-Tianjin-Hebei region in the northeast of mainland China and its neighboring Shandong, Shanxi, Henan provinces and Xinjiang areas in the west of mainland China. In addition, the concentration of NO2 was also higher in such economically developed regions as Yangtze River Delta, Pearl River Delta and Sichuan Basin, which indicates the distribution of NO2 was of universality, a different characteristic compared with the concentricity occupied by the other parameters. By contrast, O3, one of the atmospheric pollution parameters, was mainly distributed in Qinghai and Inner Mongolia in the central north of mainland China. PM2.5, AQI and PM10 experienced almost the same variation trend with time, which also happened to CO, NO2 and SO2. Whereas, O3 witnessed an opposite variation trend to CO, NO2 and SO2 with time. These cities including Baoding, Xingtai, Handan, Tangshan in Beijing-Tianjin-Hebei region, Langfang in Shandong province, Linfen in Shanxi province, haibeizhou in Qinghai, Kashi, Hetian, Aksu and Kezhou in Xinjiang should be emphatically improved. Except for the decline of the highest concentration of CO and SO2 from 2014 to 2019, this indicator of other parameters was not improved. In general, the atmospheric environment of mainland China had not been obviously improved over the six years. As a consequence, the study in this paper urges us to pay attention to the problem with the atmospheric environment of northern region of China, and also indicates cities and regions with severer atmospheric contamination, for which authentic actions should be implemented to address the corresponding problems.

